# Integrative *PTEN* Enhancer Discovery Reveals a New Model of Enhancer Organization

**DOI:** 10.1101/2023.09.20.558459

**Authors:** Christian G. Cerda-Smith, Haley M. Hutchinson, Annie Liu, Viraat Y. Goel, Corriene Sept, Holly Kim, Salvador Casaní-Galdón, Katherine G. Burkman, Christopher F. Bassil, Anders S. Hansen, Martin J. Aryee, Sarah E. Johnstone, Christine E. Eyler, Kris C. Wood

## Abstract

Enhancers possess both structural elements mediating promoter looping and functional elements mediating gene expression. Traditional models of enhancer-mediated gene regulation imply genomic overlap or immediate adjacency of these elements. We test this model by combining densely-tiled CRISPRa screening with nucleosome-resolution Region Capture Micro-C topology analysis. Using this integrated approach, we comprehensively define the *cis*-regulatory landscape for the tumor suppressor *PTEN*, identifying and validating 10 distinct enhancers and defining their 3D spatial organization. Unexpectedly, we identify several long-range functional enhancers whose promoter proximity is facilitated by chromatin loop anchors several kilobases away, and demonstrate that accounting for this spatial separation improves the computational prediction of validated enhancers. Thus, we propose a new model of enhancer organization incorporating spatial separation of essential functional and structural components.

## Main Text

Enhancers drive gene expression patterns that dictate critical cellular processes.^1^ These non-coding genomic elements transcriptionally activate gene targets on the same linear DNA molecule in dynamic and context-specific ways, influencing human physiology and disease.^1,2^ Hundreds of thousands of putative enhancers have been identified through various genome-wide biochemical and sequencing-based approaches.^3,4^ Despite these efforts, aside from a few well-characterized loci, little is known about the specific fine-scale mechanics by which enhancers physically interact with specific gene partners.^2^

Canonically, enhancers recruit and deliver transcription factors to gene promoters via physical three-dimensional (3D) looping patterns facilitating enhancer-promoter (E-P) contact.^5–8^ Such models assume that functional enhancers make direct structural contact with promoters via chromatin loops.^1,5,7^ However, existing experimental evidence lacks the resolution to provide specific insight into fine-scale E-P chromatin loop architecture.^9,10^ This limits understanding of how the organization of specific structural and functional enhancer components directly influence gene regulatory activity.

We reasoned that integrating high-resolution functional enhancer profiling and nuclear topology analyses would clarify the spatial relationship between functional enhancers and structural elements mediating promoter contact. We thus paired two methods with unparalleled resolution: 1) a recently described Region Capture Micro-C (RCMC) genome topology mapping approach permitting nucleosome-level (150bp) resolution, and 2) a high-density tiled-CRISPR activation (CRISPRa) enhancer discovery approach. We employed these methods to characterize the *cis*- regulome of the haploinsufficient tumor suppressor gene *PTEN*, as deeper understanding of its noncoding regulation could help explain its frequent loss in tumors and fuel efforts toward therapeutic restoration.^11–15^ Our RCMC study yielded the highest-resolution nuclear topology landscape over a large genomic window in human cells to date. By pairing this with a CRISPRa enhancer discovery approach, we identified and validated 10 distinct *PTEN* enhancers, including 9 previously undescribed, acting at distances up to 565 kb from the *PTEN* promoter. Our integrative approach unveiled notable heterogeneity in E-P organization. Most notably, this analysis revealed a new archetype for enhancer organization wherein significant genomic distances may separate functional enhancer elements from structural anchors mediating promoter proximity. This regulatory organization is observed at diverse gene loci and has important implications for widely used computational enhancer discovery models that assume direct promoter-enhancer contact.^16^

## Results

### Region Capture Micro-C (RCMC) Defines the Topologically Complex *PTEN* Locus

Prevailing enhancer models assume topologic contact with gene promoters (E-P contact).^1,5–8,16^ We reasoned that the topologic organization around *PTEN* would provide critical insight into the gene’s *cis*-regulatory landscape. We therefore interrogated existing published chromatin conformation capture (3C) datasets to identify likely *PTEN* enhancers, but even the highest resolution datasets failed to fully detail fine-scale chromatin contacts (**Fig. 1a-c**).^17^

**Fig. 1.**
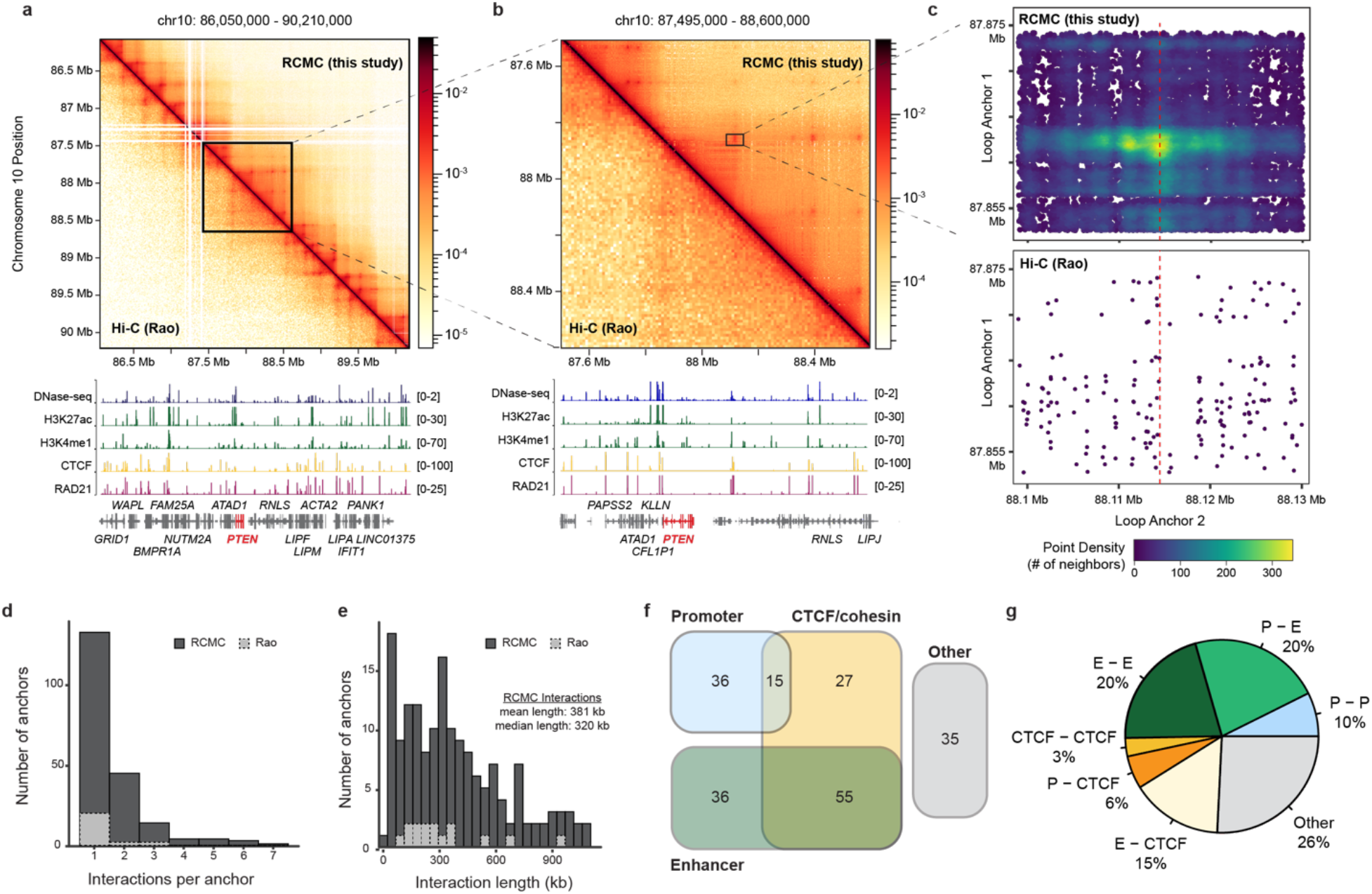
RCMC yields enhanced high-resolution contact map for large genomic window enabling detection of novel structure. Contact map comparison of HCT116 RCMC against Hi-C (Rao 2017) **(a)** scaled to *PTEN* capture window of 4.16 Mb, and **(b)** 1.105 Mb surrounding *PTEN* locus. **(c)** Scatterplot of pairwise contacts, within representative contact-containing window, generated by RCMC versus Hi-C. Density function indicates degree of overlapping points; purple represents single paired-read, yellow represents overlapping paired-reads. **(d)** Distribution of number of interactions formed by each detected anchor in RCMC vs Hi-C. **(e)** Distribution of interaction lengths detected in RCMC vs Hi-C. **(f)** Venn diagram of RCMC detected anchor categories determined by overlapping chromatin features within 5kb of anchor midpoints. Promoters defined as TSS +/- 3kb. Enhancers defined by coinciding histone marks (H3K27ac, H3K4me1) determined by ChIP-seq, not overlapping promoters. CTCF/Cohesin defined by coinciding CTCF and Rad21 binding determined by ChIP-seq. Other regions are those not overlapping any such features. **(g)** Fractions of loops classified into different categories: P-P (promoter-promoter), E-P (enhancer-promoter), E-E (enhancer-enhancer), CTCF-CTCF (CTCF/cohesin-CTCF/cohesin), P-CTCF (promoter-CTCF/cohesin), E-CTCF (enhancer-CTCF/cohesin), Other (Other-Other). For panels (a) and (b): visualization of each contact map was optimized to the technique and viewing window; here, RCMC=2kb bins and Hi-C=10kb bins. Gene annotations, DNase-seq (blue) and ChIP-seq tracks (H3K27ac=green, H3K4me1=green, CTCF=yellow, RAD21=pink) shown. *PTEN* in red.

To clarify the genomic context and three-dimensional organization of the *PTEN* locus, we employed RCMC, a recently described 3C technique.^18^ This method achieves nucleosome-resolution by combining micrococcal nuclease fragmentation, locus-specific oligo capture, and efficient deep sequencing **(Extended Data Fig. 1)**. RCMC dramatically clarified the topologic organization of our genomic window in HCT116 cells, unveiling numerous new putative regulatory contacts beyond those evident in previous datasets (**Figs. 1a-c**). Indeed, our RCMC study represented a nearly 17-fold improvement in resolution over the most resolved published HCT116 Hi-C dataset, with calculated resolutions of 150bp versus 2500bp, respectively **(Extended Data Fig. 2)**.^17,19^ To our knowledge, this resolution across a profiled genomic region of 4.16 Mb represents the highest resolution topology study for a continuous genomic region of its size in human cell lines.

This approach identified 204 unique chromatin loop anchors forming 163 unique looping contacts, with contact distances ranging from 20 - 1098 kb (mean 381 kb, median 320 kb) **(Figs. 1d-e, Extended Data Fig. 3a)**. This exceeded similar re-analysis of previously published genome-scale HCT116 Hi-C datasets, which revealed 24 unique anchors forming 15 unique contacts, with contact distances ranging from 100 - 930 kb (mean 329 kb, median 260 kb) **(Fig. 1d-e, Extended Data Fig. 3b)**.^17^ In evaluating the chromatin context of loop anchors detected by RCMC, 45% overlapped putative enhancers, 25% coincided with promoters, and 48% corresponded to CTCF- and cohesin-bound anchors (**Fig. 1f**). This represents detection of 33 enhancer-promoter, 16 promoter-promoter, and 33 enhancer-enhancer contacts (**Fig. 1g**). The region immediately flanking *PTEN* exhibited markedly complex topologic patterns, including numerous interactions extending outside of topologically associating domains (TADs), and was enriched for anchors forming chromatin loops (25% of loop anchors, participating in >53% of called loops in the profiled region, fell within the 900 kb region flanking *PTEN*) **(Figs. 1a-b, Extended Data Fig. 3a)**. Thus, our RCMC topology map provided an unparalleled view of the complex topologic relationships involving the *PTEN* locus.

### High-Resolution Tiled CRISPRa Screening Nominates *PTEN* Functional Enhancers

Topologic approaches like RCMC provide critical insight into 3D organization of a locus and may nominate candidate E-P gene interactions.^9,18,20^ However, topology alone cannot assess the functional significance of detected contacts.^2,10^ Thus, we performed a parallel analysis using flow cytometry-based, tiled CRISPRa screening to identify functional *PTEN* enhancers.^21,22^ Because the *PTEN* promoter lies at a TAD boundary and our RCMC map indicated numerous extra-TAD looping interactions, we interrogated a 5 Mb window surrounding *PTEN* to maximize the likelihood of capturing all potential *PTEN* enhancers **(Figs. 1a-b, Extended Data Fig. 3a)**. To facilitate this breadth and enrich tiling density, we targeted all putative specific sgRNA binding sites within the composite 623 DNase hypersensitive (DHS) regions compiled from 5 *PTEN* wildtype cell lines (**Fig. 2a**).^22,23^

**Fig. 2.**
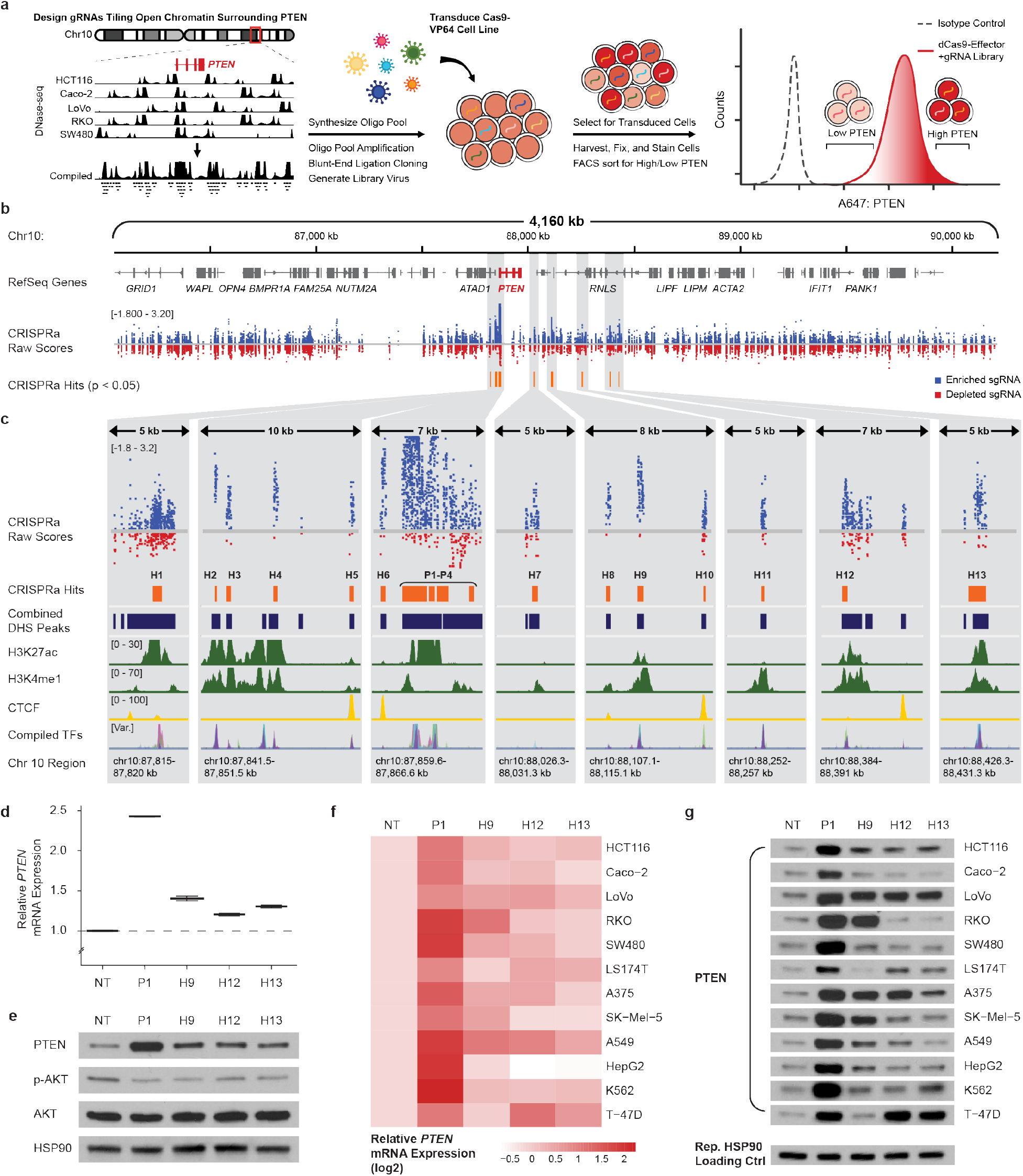
Tiled CRISPRa-based epigenomic screen in HCT116 identifies both novel and previously identified *PTEN* regulatory elements. **(a)** CRISPRa tiling screen workflow. **(b)** CRISPR-SURF analysis comparing sgRNA enrichment/depletion between “PTEN-High” and control “All” populations. **(c)** Zoom-ins on select significant regions located upstream, within promoter, and downstream of *PTEN* TSS with coinciding biochemical marks. Single sgRNA validation of downstream hit regions (“H9”, “H12”, “H13”) **(d)** via qRT-PCR (n=4 technical replicates for each of n=3 biological replicates; mean and standard error depicted), and **(e)** western blot in HCT116 (representative of n=3 biological replicates). Single sgRNA validation of downstream hit regions (“H9”, “H12”, “H13”) across cell lines **(f)** via qRT-PCR (visualized as heatmap), and **(g)** via western blot. FACS = fluorescence-activated cell sorting. For panels (b) and (c), squares represent individual sgRNAs (blue=log fold change > 0, red=log fold change < 0 in the “PTEN-High” versus control “All” populations), orange bars represent called significant hit regions, dark blue bars represent open chromatin regions defined by DNase-seq, green peaks represent enhancer-associated histone marks (H3K27ac, H3K4me1) determined by ChIP-seq, yellow peaks represent CTCF binding determined by ChIP-seq, and multi-coloured peak track represents various TF binding determined by ChIP-seq (POLR2AphosphoS5=pink, JUND=peach, YY1=lime green, FOSL1=teal, SP1=violet, TCF7L2=light blue, SRF=navy, CBX3=purple). For panels (d) - (g): NT=non-targeting control, P1=promoter positive control

Using HCT116 cells stably expressing nuclease-deactivated Cas9 fused to two VP64 transcriptional activator moieties (VP64-dCas9-VP64), we performed CRISPRa screening with our high-density tiled lentiviral library, flow-sorting populations by PTEN expression (PTEN High, PTEN low, and PTEN all [control]), and subsequently analyzed sgRNA enrichment via CRISPR-SURF (**Fig. 2a-b**).^21,24^

Comparison of the “PTEN High” population with the “PTEN All” control nominated 13 discrete hit regions as candidate *PTEN* enhancers (“H1”–“H13”), and called 4 positive controls spanning the *PTEN* promoter (“P1”–“P4”) (**Fig. 2b, Extended Data Table 1**). Hits ranged from 45 kb upstream to 565 kb downstream of the transcription start site (TSS) (**Fig. 2b-c, Extended Data Table 1**). All identified candidate enhancers fell within HCT116-specific DHS regions despite screening significant genomic space outside of these territories, underscoring the specificity of this approach. Moreover, this method’s high resolution pinpointed specific candidate enhancer sub-regions within broader DHS domains due to dense sgRNA tiling and the DHS-agnostic analysis pipeline (**Fig. 2c**). Importantly, our screen re-identified the only two previously described *PTEN* enhancers, labeled “P1” and “H12” here (P1 lies in the region we refer to as promoter).^25,26^ To our knowledge, this is the first report of the remaining 12 *PTEN* candidate enhancers (“H1”-“H11”, “H13”). As expected for functionally active enhancers, hits overlapped with canonical enhancer histone marks (H3K27Ac, H3K4Me1) and co-localized with structurally and functionally important transcription factors (CTCF, YY1, POLR2AphosphoS5, JUND, FOSL1, SP1, TCF7L2, SRF, CBX3) (**Fig. 2c**).^1,27^ In validation experiments testing 13 non-promoter hits, functional activation of 10 significantly increased PTEN expression (**Figs. 2d-e, Extended Data Fig. 4**). Further interrogation of the strongest long-range (>25 kb from TSS) validated enhancers demonstrated corresponding repression of canonical downstream AKT phosphorylation, suggesting that individual enhancers can functionally modulate PTEN expression to meaningful degrees (**Fig. 2e**).^28^

Enhancers exhibit tissue variability; it is unclear whether enhancers identified in one context function in other settings.^6,8,23,29^ Thus, we evaluated the strongest long-range enhancers from our HCT116 study across various cell lines. While *PTEN* promoter activation increased expression in 12 cell lines from 6 different tissues, the HCT116-identified enhancers demonstrated cell-type variability in modulating expression (**Figs. 2f-g, Extended Data Fig. 5**). The majority of cell lines upregulated PTEN via at least 1 of the HCT116-identified enhancers, with perturbation of each enhancer causing significant expression changes in at least 3/12. Thus, while we observe expected cell-type variability, these enhancers function as regulators in multiple cellular contexts.

### Integrated Enhancer Discovery Analysis Reveals Spatially-Distinct Functional and Structural Components

Enhancers are widely thought to 1) functionally regulate gene expression, and 2) make topologic contact with/near gene promoters (E-P contact).^1,2,6,7^ It is commonly assumed that the functional and topologic components are genomically overlapping or directly adjacent. However, accurate depictions of enhancer organization require dedicated high-resolution assays profiling both chromatin looping and functional enhancers, which had not previously been performed together.

We cross-referenced our HCT116 RCMC topologic map with our screen’s functionally validated enhancers to assess the spatial relationship between functional enhancers and structural anchor points mediating promoter proximity (**Fig. 3a**). Surprisingly, only 1/5 functionally validated long-range enhancers (“H13”) directly overlapped with a chromatin loop anchor (**Fig. 3b-c**); moreover, none coincided with CTCF binding, which is commonly described as driving long-range enhancer loops (**Fig. 3b-c**).^30,31^ The average distance from long-range functional enhancers to the nearest CTCF/cohesin coinciding anchor was 39.69 kb (median 39.63 kb, range 3.39-86.23 kb), while the average distance from the nearest general RCMC anchor point was 11.15 kb (median 3.39 kb, range 0.37-44.23 kb) (**Fig. 3b-c**). In fact, 4 of the strongest functional enhancers (“H7”, “H9”, “H11”, “H12”) coincided with neither a called structural anchor nor CTCF/cohesin (**Fig. 3b-d**). Further, for the two such long-range non-anchor-overlapping functional hits with closest proximity to a loop anchor (“H9”, “H12”), the effect of CRISPRa-based activation of the nearest anchor points (“H10”, “A12”, respectively) alone on PTEN expression did not reach statistical significance (**Extended Data Fig. 4b**). This highlights the specificity of CRISPRa for functional enhancer identification and underscores the genomic separation of functional enhancer components from the structural chromatin loop anchors facilitating promoter proximity.

**Fig. 3.**
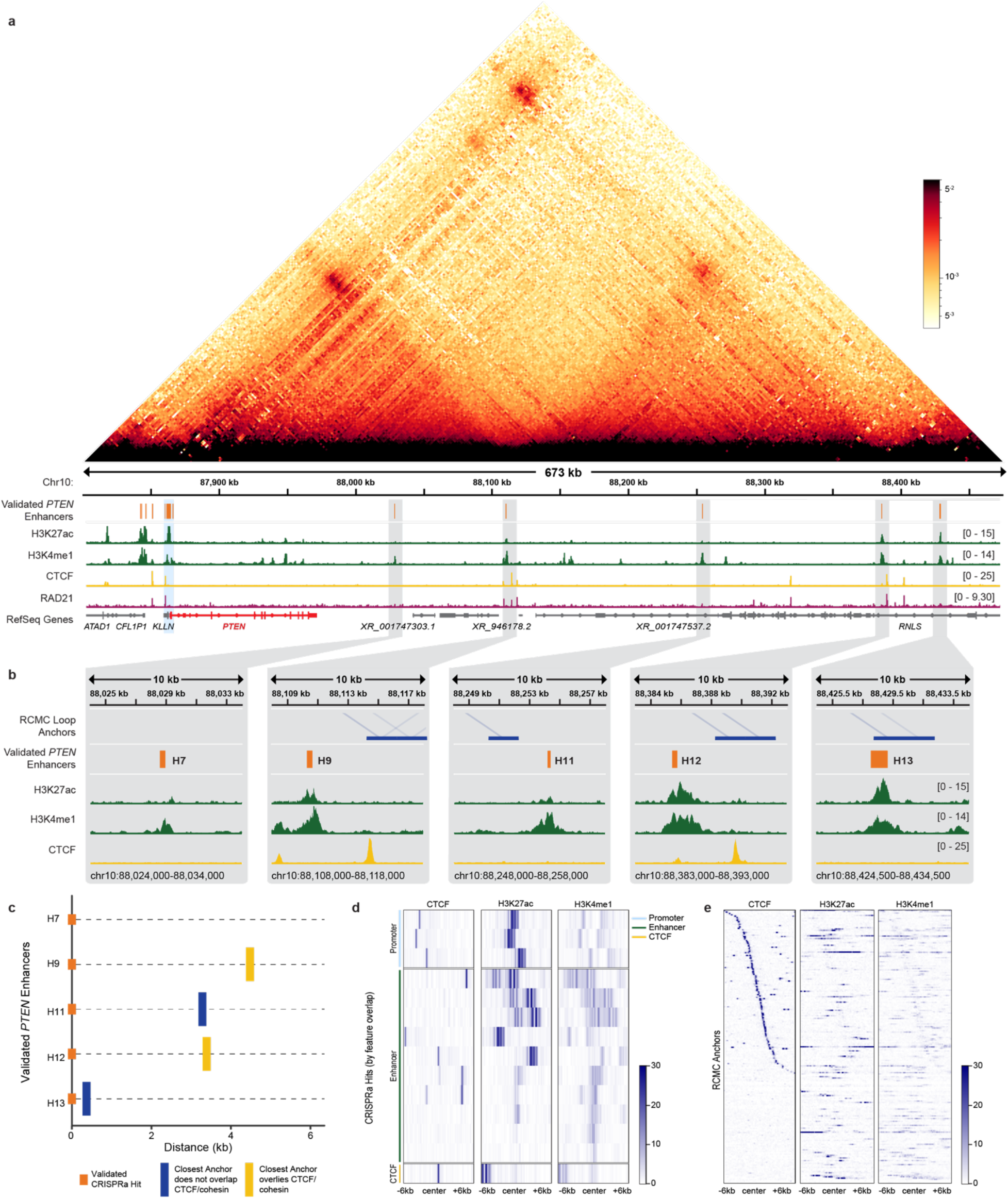
Integration of RCMC topological and CRISPRa functional platforms reveals resolution of enhancer structural and functional components. **(a)** Overlay of RCMC contact map and functionally validated enhancer track from tiled CRISPRa screen (orange) in 673 kb window surrounding *PTEN* locus. **(b)** Zoom-ins of regions containing functionally validated long-range (>25kb) *PTEN* enhancers (orange). Called loop anchors are shown (blue). **(c)** Plot visualizing functionally validated long-range (>25kb) *PTEN* enhancers (orange) and absolute distance to nearest loop anchor (blue, or yellow if anchor overlaps CTCF/cohesin peak). Enhancer tracks without displayed anchor don’t have one detectable within 25kb of the functional enhancer. **(d)** Heatmap depicting localization of designated feature (x-axis) relative to functionally validated *PTEN* regulatory elements (y-axis). *PTEN* regulatory elements subdivided based upon underlying biochemical features (promoter=light blue, enhancer=green, CTCF-bound=yellow) **(e)** Heatmap depicting localization of designated feature (x-axis) relative to RCMC-determined anchors (y-axis) across 4.16 Mb window (y-axis). For panels (a) through (c): Gene annotations shown, *PTEN* gene in red. *PTEN* promoter highlighted in light blue. Functionally validated long-range (>25 kb) *PTEN* enhancers highlighted in gray. Functionally validated enhancers=orange bars, ChIP-seq tracks (H3K27ac=green, H3K4me1=green, CTCF=yellow, RAD21=pink) are shown.

To assess the generalizability of our findings, we queried the genomic relationship of RCMC chromatin loop anchors and putative functional enhancer elements across the entire 4.16 Mb RCMC-profiled region. Enhancer markers (H3K27Ac, H3K4Me1) directly overlapped with RCMC promoter-linked chromatin anchors in 31.4% of cases, consistent with direct E-P contact (**Fig. 3e**). However, in 68.6% of cases, putative enhancer markers did not directly overlap topologic promoter-proximal anchors. In fact, the average genomic separation of putative functional and structural components across our 4.16 Mb window was 7.32 kb (median 3.48 kb) (**Fig. 3e**). By integrating high-resolution contact mapping with high-density functional profiling methods, we observe that a significant fraction of functional enhancers are spatially separated from the genomic regions facilitating long-range promoter proximity.

### Enhancer Prediction Model Adjustments Correct for Spatial Gaps Between Topologic and Functional Enhancer Components

Current notions of enhancer function build on the recognition that long-distance chromatin looping facilitates enhancer contact with gene promoters.^1,6–8^ Expanding upon this, the Activity-by-Contact (ABC) computational model facilitates integration of biochemical enhancer annotations with topologic features to predict E-P pairs on a genome-wide scale.^16,32^ We were eager to compare our *PTEN* enhancers with those predicted by ABC. We noted, however, that ABC predicted few *PTEN* enhancers in HCT116 using published Hi-C maps. Indeed, these analyses predicted only 4/10 functionally validated enhancers (“H2”, “H4”-“H6”), even when liberalizing ABC to accept both H3K27Ac and H3K4Me1 as putative enhancer markers. Notably, the predicted enhancers all resided upstream and within 25 kb of the TSS, missing the long-range and strongest enhancers. We first suspected this limitation related to limited Hi-C data density restricting the ability to predict subtle E-P contacts. Given our dataset’s improved resolution and read density, we re-ran the ABC model using our RCMC data. Here, too, ABC was unable to predict the long-range (>25 kb) screen-identified and/or previously reported enhancers (**Fig. 4a-b**).

**Fig. 4.**
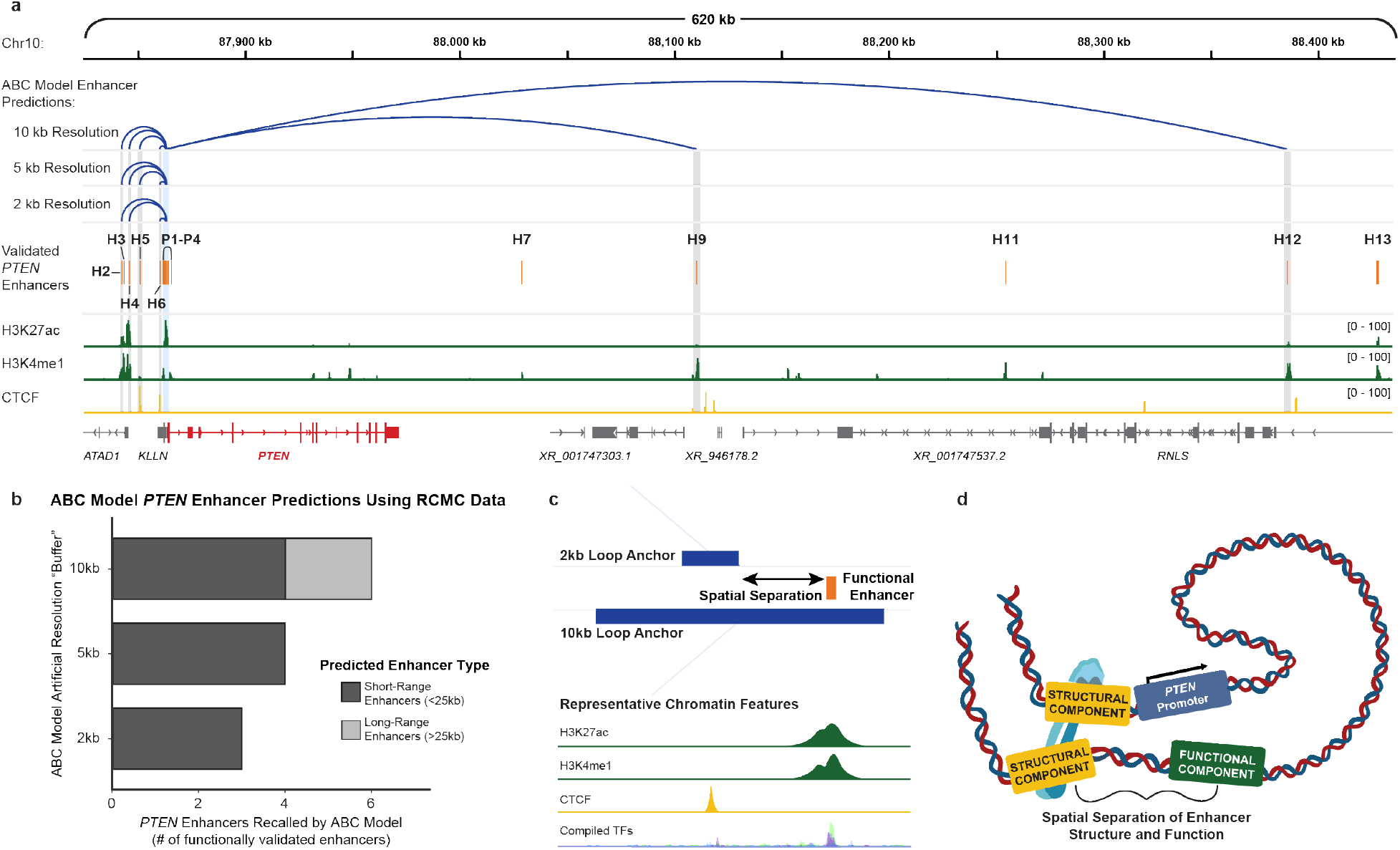
Enhancer prediction model fails at high resolutions due to spatial separation between topologic and functional enhancer components. **(a)** ABC model predictions (blue loops) informed by RCMC data at 10kb, 5kb, and 2kb resolutions versus functionally validated *PTEN* enhancers (orange). The *PTEN* promoter region is highlighted in light blue. Functionally validated *PTEN* enhancers are highlighted in gray. Gene annotations are shown, *PTEN* gene in red. **(b)** Plot showing ability of ABC model informed by RCMC data to recall validated *PTEN* enhancers at 10kb, 5kb, and 2kb resolutions. Short-range enhancers (<25kb)=dark gray, long-range enhancers(>25kb)=light gray. **(c)** Schematic illustrating effect of spatial separation of enhancer functional and structural elements on enhancer prediction algorithms utilizing high-resolution datasets. **(d)** Model of enhancer organization illustrating potential for spatial separation of enhancer functional components from structural components mediating promotor-proximity. For panels (a) and (c): functionally validated enhancers=orange, ABC-predicted enhancers=dark blue lines/loops, RCMC-detected loop anchors=dark blue bars, green peaks represent enhancer-associated histone marks (H3K27ac, H3K4me1) as determined by ChIP-seq, and yellow peaks represent CTCF binding as determined by ChIP-seq.

We hypothesized that ABC’s failure here derived from spatial gaps separating functional elements from structural components that mediate looping (**Fig. 4c**). Indeed, artificially limiting RCMC resolution to facilitate “buffered” overlap of detected loop anchors with functional enhancers increased ABC’s ability to recall our study’s functionally validated enhancers. Limiting resolution to 10 kb enabled prediction of 6/10 enhancers including two long-range enhancers (“H9”, “H12”). As we decreased this artificial resolution “buffer” from 10 kb, to 5 kb, to 2kb, ability to predict functionally validated enhancers diminished (6 recalled enhancers [2 long-range], 4 recalled enhancers [0 long-range], and 3 recalled enhancers [0 long-range], respectively) (**Fig. 4a-c**). In other words, accurate prediction of functional enhancers using new high-resolution topology datasets derived from RCMC or Micro-C requires adjustments (e.g., incorporation of a several kilobase “buffer” around RCMC loops) to account for possible spatial separation between functional enhancers and corresponding structural elements (**Fig. 4c**). Alongside the proliferation of high-resolution topologic datasets, ongoing efforts to improve computational enhancer prediction models would benefit from incorporating the possibility of spatial separation between functional enhancer elements and their promoter-contacting chromatin loops.

By integrating unparalleled RCMC topologic resolution with functional enhancers nominated via high-density CRISPRa screening, we reveal that functionally active enhancer regions can be physically discrete from structural elements mediating promoter proximity. These non-adjacent functional and structural units are evident within the *PTEN* locus as well as across a variety of genes within a >4 Mb genomic window. Based on these findings we propose a revised model of enhancer structure and function whereby a broad gene regulatory domain possesses both functionally active elements and structurally critical components which may not exhibit genomic adjacency or overlap (**Fig. 4d**). Our revised model of enhancer organization has important implications. It clarifies canonical E-P interaction models, which often assume that chromatin loops spatially correspond with functionally active enhancer regions.^1,7,33^ Our data largely confirm the anticipated proximity of enhancers with promoters, but clarify that many “E-P loops” are strengthened by anchors that are actually several kilobases away from their paired functional elements.

Previously, methodologic limitations prevented systematic detection of spatio-functional discordance between the location of functional enhancers and promoter-contacting structural loop anchors, though this arrangement has been theorized.^34,35^ We surmise that chromatin looping facilitated by these spatially-resolved loop anchors could draw functional enhancers close enough to promoters for short-range diffusion of activating factors that facilitate transcription in a looping-then-diffusion model of transcription factor activation.^36,37^ Alternatively, but not mutually exclusively, strong loops provided by structural elements may suffice to draw spatially separated functional enhancers close enough to promoters to permit “bridging” or “direct extension” of transcription factors and activators in large complexes.^35^

Direct E-P contact may be unnecessary for promoter activation by functional enhancers. Given the spatial heterogeneity by which functional enhancers interact with promoters across our profiled locus, we propose that computational enhancer prediction methods will benefit from integrating the potential spatial discrepancy between functional and topologic elements, particularly as technology continues to produce higher resolution datasets capable of resolving finer-scale structures. For example, we demonstrate that ABC could easily be adapted to account for spatially-distinct structural and functional components evident in high-resolution topology datasets like RCMC.^16^ Therefore, we anticipate such methods will continue to drive large-scale enhancer prediction efforts after accounting for this new model of enhancer structure.

Other surprising aspects of the *PTEN* topologic landscape deserve comment. We observed multiple clustered loop anchors in regions where Hi-C had detected only single loops, nominating short-range chromatin loop redundancy as one mechanism by which E-P interactions might resist perturbation, as observed elsewhere.^38^ In addition to observed CTCF-dependent looping, many CTCF-independent loops promoting enhancer-promoter proximity were identified, consistent with notions of alternative mechanisms of chromatin organization including those driven by non-CTCF looping mediators, microcompartments, or proximity.^18,20,39–41^ We identified numerous functionally active enhancers >100 kb from the TSS, adding to the catalogue of known enhancers acting at very long distances. Notably, these long-range E-P contacts fall within the broader *PTEN*- containing TAD, consistent with reports of human embryonic stem cells demonstrating intra-TAD loops, and in Drosophila where intra-TAD “tethers” facilitate coordinated multigene regulation.^42,43^ Interestingly, we observed evidence of contacts between several validated long-range enhancers suggesting a potential role for previously described multiway contacts in the regulation of *PTEN*, warranting further investigation.^44^ Though beyond the scope of this study, while all identified long-range *PTEN* enhancers fell within the *PTEN*-containing TAD, RCMC enabled detection of looping interactions between these enhancer elements and non-functional regions extending beyond TAD boundaries, supporting recent descriptions of TAD permeability and allowing for potential cross-TAD structural regulation of enhancers.^45^

Our *PTEN*-specific findings have far-reaching implications, as non-mutational PTEN repression impacts disease.^11–13,46^ As a haploinsufficient tumor suppressor, even small decreases in wild type PTEN expression correlate with increased cancer susceptibilities and impact cancer cell proliferation, apoptosis, drug sensitivity, and immune interactions.^11,14,46–48^ Many patients with clinically diagnosed PTEN Hamartoma Tumor Syndromes exhibit non-mutational mechanisms of PTEN repression, which could include aberrant *cis*-regulation of *PTEN* transcription.^12,49^ By characterizing the functional and structural context of 9 novel and 2 previously reported *PTEN* enhancers, and validating their activity across various contexts, we provide a new understanding of *PTEN* regulation with potential diagnostic or therapeutic implications.^15,25,26,50^

Methodologically, our combined functional and topologic approach provides an objective and reproducible framework for future enhancer discovery. First, the locus specificity of the functional and topologic components allows high-density information at relatively low cost, augmenting practical feasibility. Further, our observation of promoter-contacting loops far exceeding the number of identified functional enhancers reaffirms the inadequacy of topology alone for nominating enhancers.^2,10^ Additionally, high-density tiled CRISPRa identifies regions with regulatory potential including “poised” enhancers or functionally redundant enhancers, compared to CRISPRi which assesses the necessity of enhancers for baseline expression.^1,2,21,22^ Countering one criticism of CRISPRa as prone to “off-target” hits, we observed that looping to the promoter alone was insufficient for CRISPRa-mediated PTEN expression, and hits instead appeared specific to putative enhancers possessing functional biochemical annotations (e.g., H3K27Ac, H3K4Me1). Further, despite querying a large genomic region, our results indicate that enhancers with strong *PTEN* activation potential all localized to the TAD containing *PTEN* and all coincided with cell line-specific accessible DNA. As we found that several long-range enhancers existed well beyond 100 kb from the TSS, but within the same TAD as their gene partner, topologic data may provide valuable guidance in tiling library design versus a strict distance metric.

In summary, we report an advanced, integrative platform for enhancer discovery that combines screen-based functional perturbations with detailed topologic data to comprehensively define the structural-functional *cis*-regulome of specific genes. Our analysis of the *PTEN* locus allowed us to annotate and validate many new enhancer elements. Most significantly, we reveal new patterns characterizing the spatial relationship between functional enhancers and the chromatin loops that facilitate promoter proximity. We posit a new model for enhancer organization whereby a broad enhancer domain possesses both functionally active and structurally critical components that, in contrast to their previously assumed genomic overlap or adjacency, can be separated by multi-kilobase distances.

## Methods

### Cell Culture

A375, A549, Caco-2, HCT116, HepG2, HEK293T, K562, LoVo, LS174-T, RKO, SK-Mel-5, SW480, and T-47D cell lines were purchased from the American Tissue Collection Center (ATCC) via the Duke University Cell Culture Facility (Duke CCF). All cell lines were authenticated by short tandem repeat (STR) profiling and verified mycoplasma free using the MycoAlert PLUS Mycoplasma Detection kit (Lonza) prior to conducting experiments. All cells were maintained at 37°C at 5% CO_2_ and cultured in the respective media outlined in (Extended Data Table 2).

### Region Capture Micro-C (RCMC)

We performed the protocol as preliminarily described, with some modifications.^18^ Briefly, 2.5×10^7^ HCT116 cells were crosslinked with DSG and formaldehyde, in biologic duplicate. The chromatin was digested to mononucleosomes utilizing a micrococcal nuclease (MNase) digestion first optimized via titration, followed by end-repair with biotin labeling, and proximity ligation. Samples were then reverse-crosslinked, size-selected for dinucleosome fragments, and purified for ligated fragments via streptavidin pulldown. The resultant DNA was processed into Micro-C sequencing libraries via adapter ligation and PCR. We designed a custom 120-mer oligo capture pool tiling a 4.16Mb window surrounding the *PTEN* locus; the capture pool was designed at 2x coverage of the genomic window with filtering for highly repetitive regions, ultimately yielding a pool of 60,570 oligos that covered 90.16% of the region. The capture pool was synthesized and purchased as a Custom Target Enrichment Panel from Twist Bioscience. Oligo-mediated locus-specific capture and RCMC sequencing library preparation were then performed using our custom oligo capture pool following the Twist Fast Hybridization Target Enrichment Protocol. The library was sequenced on the NovaSeq 6000 platform (Illumina) yielding 814M 101PE reads; importantly, increasing the read length from 50bp to 101bp enabled efficient capture of the ligation junction in nearly 50% of reads, allowing for enhanced resolution.

### Analysis and Visualization of Topologic Data

Analyses of RCMC and Hi-C datasets were performed as preliminarily described.^18^ Briefly, paired-end reads were aligned to the UCSC hg38 genome using bowtie2 (v2.3.5.1), and aligned read mates were subsequently processed with pairtools (v0.3.0) to remove PCR duplicates and non-uniquely mapped reads, index reads, and filter to retain only reads with both mates within the captured region (**Extended Data Table 3**).^17,51,52^ Read counts were binned across the genome for 50bp bins to coarser resolutions using cooler (v0.8.11).^53^ Resulting contact matrices were balanced using iterative correction and eigendecomposition (ICE) within the captured region.^54^ The balanced contact matrix was visualized for preliminary analysis using HiGlass (v0.8.0) and for figure generation using cooltools (v0.5.0).^55,56^ Annotations were determined via CoolBox (v0.3.3).^57^

### Resolution Determination

Resolution for both RCMC and Hi-C datasets was determined as previously described.^19^ Briefly, for increasing bin sizes (50, 100, 150, 200, 250, 300, 400, 500, 600, 700, 800, 900, 1000, 1250, 1500, 1750, 2000, 2500, 3000, 4000, 5000, 6000, 7000, 8000, 9000, 10000), the number of contacts in each bin across the 4.16Mb capture locus was counted until the number of bins with >1000 contacts was greater than 80% of the total number of bins. For the RCMC dataset this value was 150bp (which approximates nucleosome fragment size enabled by MNase digestion), and for the Hi-C dataset it was 2500bp.

### Loop Calling

For both RCMC and Hi-C datasets, chromatin loops were detected using Mustache (v1.2.4) using optimized parameters as described.^58^ For RCMC, loop calls were optimized at 2 kb data resolution with a sigma0 of 2.1, sparsity threshold of 0.7, and q-value threshold of 0.075 **(Extended Data Fig. 2)**. For Hi-C, loop calls were optimized at 10 kb data resolution with a sigma0 of 1.6, sparsity threshold of 0.88, and q-value threshold of 0.1 **(Extended Data Fig. 2)**.

### Loop Anchor Classification

RCMC loop anchors, defined as the midpoint of the Mustache-called loop anchor ±5 kb, were classified as promoter, enhancer, and/or CTCF/cohesin-bound (**Fig. 1f**) as follows. Anchors were categorized as promoters if they overlapped the consensus TSS locations in the hg38 UCSC RefGene track (version 2017-03-08) ±3 kb as determined using bedtools (v2.30.0) intersect.^59^ Non-promoter anchors were categorized as enhancers if they overlapped called peaks from either H3K27ac (ENCODE accession ENCFF899XEF) or H3K4me1 (ENCODE accession ENCFF931YSQ). Anchors were classified as CTCF/cohesin-bound regions if they overlapped called peaks from CTCF (ENCODE accession ENCFF970MXD) and RAD21 (ENCODE accession ENCFF391AAM). Putative enhancer regions were determined by combining H3K27ac and H3K4me1 narrowPeak calls from ENCODE and collapsing the resulting file using bedtools (v2.30.0) merge. CTCF/cohesin-bound regions were determined using bedtools intersect to return the overlaps between CTCF and RAD21 narrowPeak calls obtained from ENCODE. Anchors overlapping both H3K27ac/H3K4me1 and CTCF/cohesin-bound sites were classified as enhancers. Anchors that did not overlap promoters, enhancers, or CTCF/cohesin-bound sites were classified as “Other”. Anchors were then aligned with their looping partners to generate loop classifications (**Fig. 1g**).

### Heatmap Generation

Heatmaps detailing CTCF, H3K27ac, and H3K4me1 chromatin features surrounding validated *PTEN* enhancers and all RCMC loop anchors were generated using deeptools (v3.5.1) computeMatrix followed by plotHeatmap in a region ±6 kb around the center of each site.^60^ For each comparison, bigWig files from ChIP-seq experiments were obtained from ENCODE. Loop anchors in **Fig. 3e** were manually sorted based on the relative distance to the nearest CTCF peak as determined using bedtools (v2.30.0) closest.

### Plasmids

We generated the VP64-dCas9-VP64 (pLV-hUbC-dSpCas9-2xVP64-T2A-PuroR) plasmid used in the tiled CRISPRa screen and subsequent validations by replacing the blasticidin-resistance cassette of pLV-hUbC-dSpCas9-2xVP64-T2A-BSD (Addgene 162333) with a puromycin resistance cassette. Briefly, the PuroR-WPRE cassette from pLV-U6-gRNA-hUbC-mCherry-P2A-PuroR (a generous gift from the laboratory of Dr. Charles Gersbach) was PCR amplified (primers in **Extended Data Table 4**) and cloned into KpnI(NEB)-digested pLV-hUbC-dSpCas9-2xVP64-T2A-BSD via blunt-end ligation cloning (NEB). pLV-U6-gRNA-hUbC-DsRed-P2A-BSR (Addgene 83919), pMD2.G (Addgene 12259), and psPAX2 (Addgene 12260) were obtained through Addgene.

### Generation of dCas9-2xVP64 Cell Lines

A375, A549, Caco-2, HCT116, HepG2, K562, LoVo, LS174-T, RKO, SK-Mel-5, SW480, and T-47D cell lines were transduced with lentiviral VP64-dCas9-VP64 vector (pLV-hUbC-dSpCas9-2xVP64-T2A-PuroR) in 6-well plates containing a final concentration of 8ug/mL polybrene in respective cell media. Media was replaced 24h post-transduction. 48h post-transduction, transduced cells were selected with puromycin for 5 days.

### Lentivirus Production and Titering

Transfection reagents were prepared in Opti-MEM reduced serum medium (Gibco) with appropriate scaling for culture surface area, according to manufacturer instructions. For VP64-dCas9-VP64 and individual sgRNA lentiviruses, for each well in a 6-well plate, ∼80% confluent HEK293T cells were transfected with 3 µg of the respective expression plasmid, 1.39 µg of psPAX2, and 0.91 µg of pMD2.G using 15.95 µL of Lipofectamine 2000 supplemented with 17.54 µL of PLUS Reagent. For the sgRNA lentiviral library, ∼80% confluent HEK293T cells were transfected in a T-225 Flask with X µg of the plasmid library, 32.4 µg of psPAX2, and 21.2 µg of pMD2.G using 374 µL of Lipofectamine 2000 supplemented with 17.54 µL of PLUS Reagent. After 6 hours following lipofection, the transfection media was replaced with fresh pre-warmed harvest media (HEK293T media supplemented with 25% FBS). After 48 hours, the viral supernatant was collected, filtered using a 0.45 µm PES filter, and either used directly or stored at −80 °C for future use.

The titer of the lentiviral sgRNA library was determined by flow cytometry. Aliquots of 3×10^6^ HCT116 cells expressing VP64-dCas9-VP64 were seeded with varying volumes of library lentivirus in 15cm dishes containing a final concentration of 8ug/mL polybrene. Transduction media was replaced with fresh media 24h later. Cells were harvested 96h post-transduction and the levels of DsRed in each sample were measured to determine the percent transduction.

### sgRNA Library Design and Cloning

To interrogate the *cis*-regulatory landscape of *PTEN* we designed a custom sgRNA library fully tiling the DNase I hypersensitive (DHS) regions within a 5Mb window surrounding the *PTEN* locus. We merged DHS sites across 5 *PTEN* wildtype colorectal cancer cell lines characterized by ENCODE (Caco-2, HCT116, LoVo, RKO, and SW480, see **Supplementary Table 1**), yielding 623 composite DHS regions, to account for cell line variability and to generate a more comprehensive view of chromatin accessibility across cancers from this tissue type. Notably, focusing on DHS regions enabled us to enrich our targeted genomic space for that most likely to be biochemically active, efficiently sample a relatively large genomic window without our library becoming prohibitively large, and achieve a high tiling density of our informed regions. GuideScan was used to filter all possible sgRNAs tiling these regions for those with a specificity score >0.2 to mitigate off-target effects, yielding a library of 16,734 sgRNAs (14,798 targeting sgRNAs plus 1936 non-targeting negative controls)^61,62^. The targeting sgRNAs directly tiled a total 256,089bp of genomic space, providing a high average tiling density of ∼1 sgRNA per 17.3bp across the composite DHS regions within our window. Our oligo pool was synthesized by Twist Bioscience and cloned into the gRNA-hUbC-DsRedExpress2-BlastR backbone using blunt-end ligation (NEB) as previously described, with minor modifications.^63^ Lentiviral sgRNA library was produced and titered as described above.

### Lentiviral sgRNA FACS Screening

HCT116 cells with stable expression of VP64-dCas9-VP64 were seeded into 15cm tissue culture dishes with 8 ug/mL polybrene and transduced with the titered lentiviral sgRNA library at a low multiplicity of infection ∼0.3 (statistically ensuring that most cells harbor no more than 1 sgRNA) to achieve greater than 1,000x coverage of the sgRNA library, in biologic triplicate. 24h post-transduction, media was replaced with fresh media containing 15 ug/mL Blasticidin and cells were selected for 96 hours. Cells were passaged and maintained above 1,000x coverage for 7 days post-transduction to allow for stable sgRNA expression and VP64-dCas9-VP64-mediated activation of target regions. Replicates were harvested, fixed and permeabilized using the eBioscience Intracellular Staining Buffer Set (Thermo) according to manufacturer recommendations, with minor modifications. Fixed and permeabilized cells were incubated overnight at 4 °C, rotating with a A647-conjugated PTEN antibody (Novus Bioscience) at 1 mL per 1.2E8 cells. For each replicate, stained cells were then sorted by FACS, collecting the top and bottom 10% of PTEN expressing cells on the basis of A647 signal; cells were collected in FBS-coated tubes, maintaining greater than 300x coverage per high and low population. An ungated control ALL sample was also collected for each replicate.

### Screen Processing and Data Analysis

Immediately following collection, samples were individually subjected to reverse-crosslinking and genomic DNA extraction using the Arcturus PicoPure DNA Extraction Kit (Thermo), according to manufacturer recommendations. sgRNA libraries were recovered from gDNA via PCR amplification using NEBNext Ultra II Q5 Master Mix (New England BioLabs) according to manufacturer instructions and using custom primers (**Extended Data Table 4**), as previously described.^22^

Amplified libraries were purified using SPRIselect beads (Beckman Coulter) employing right-sided selection of 0.8x then to 1.2x the original volume. Each DNA sample was quantified using the Quant-iT dsDNA Broad Range Assay Kit (Thermo). Samples were pooled and sequenced on a NextSeq 500 (Illumina) with 20bp single-end sequencing using custom read and index primers (**Extended Data Table 4**).

Raw sequencing reads were trimmed and processed, and analysis of enrichment and depletion metrics comparing the top 10%, bottom 10% and control ALL sorted populations were performed using the CRISPR-SURF software analysis pipeline under default settings.^24^

### Individual gRNA Cloning and Validation

The protospacers from the top enriched sgRNAs found in each region were ordered as oligonucleotides from IDT (**Supplementary Table 2**) and cloned into the Esp3I-digested lentiviral sgRNA expression vector (pLV-U6-gRNA-hUbC-DsRed-P2A-BSR) via T4 ligation. VP64-dCas9-VP64 expressing cells were individually transduced in biologic triplicate with equal volumes of each lentiviral sgRNA in 6-well plates containing a final concentration of 8ug/mL polybrene in respective cell media. 24h post-transduction, media was replaced with fresh media containing 15 ug/mL Blasticidin and cells were selected for 96 hours. For mRNA-level qRT-PCR validations, samples were harvested 6 days post-transduction. For protein-level western blot validations, samples were harvested 7 days post-transduction.

### qRT PCR

For all validations, experiments were performed in biologic triplicate. After washing with ice-cold PBS, lysate from ∼1×10^6^ cells in the well of a 6-well plate was collected in 300 µL of DNA/RNA Shield (Zymo). RNA was isolated using the Quick-RNA Miniprep Plus Kit (Zymo, R1058), quantified using the RNA Quant-IT RNA Broad Range kit (Thermofisher, Q10213), and cDNA was generated from 500 ng of RNA input using the iScript cDNA Synthesis kit (BioRad, #1708891), according to manufacturer instructions. For each biologic replicate, qRT-PCR was carried out in technical quadruplicate using the TaqMan assay (Applied Biosystems) and CFX384 Touch Real-Time PCR Detection System (Bio-Rad), according to manufacturer recommendations. The results are expressed as log_2_ fold-increase mRNA expression of PTEN normalized to TBP expression by the ΔΔCt method. Specific TaqMan gene expression assay IDs were as follows (**Extended Data Table 4**): PTEN (Taqman Probe PTEN: Hs02621230_s1 FAM-MGB) and internal housekeeping gene control (Taqman Probe TBP: Hs00427620_m1 VIC-MGB).

### Western Blotting

For all validations, experiments were performed in biologic triplicate. After washing with ice-cold PBS, whole cell lysates were obtained from ∼80% confluent cells in individual wells of 6-well dishes using RIPA lysis buffer (Sigma Aldrich, R0278-500ML) supplemented with Protease and Phosphatase Inhibitor Mini Tablets (Thermofisher) and cell scraping. Collected lysates were rotated at 4°C for 15 minutes then cleared via centrifugation at 14,000g for 10 min at 4°C. The protein was quantified using the Bio-Rad DC Protein Assay Kit (Biorad), and combined with 4X Laemmli Sample Buffer. 20-25ug of protein lysate was loaded per well and separated on a 4–20% Mini-PROTEAN® TGX Stain-Free™ Protein Gel (Bio-Rad). Proteins were transferred onto an activated PVDF membrane using the Trans-Blot Turbo Transfer system (Bio-Rad). Membranes were blocked in 5% milk in PBS-T for 1h at room temperature before incubation with primary antibodies overnight at 4°C. Membranes were washed 3 times with PBS-T before incubation with secondary antibody diluted with 5% milk in PBS-T for 1h at room temperature. After washing 3 times in PBS-T, the membranes were developed using film.

The following antibodies were used for Immunoblot analysis (**Extended Data Table 5**): Purified Mouse Anti-PTEN (BD Bioscience, Cat# 559600), Phospho-Akt (Ser473) (Cell Signaling Technology, #9271), HSP90 (C45G5) Rabbit mAb (Cell Signaling Technology, #4877), Akt (pan) (C67E7) Rabbit mAb (Cell Signaling Technology, #4691), Monoclonal ANTI-FLAG® M2 antibody produced in mouse (Sigma, F1804-200UG). All antibodies were diluted in 5% milk in PBS-T with the exception of p-AKT, which was diluted in 5% BSA in PBS-T. The PTEN antibody was prepared at a 1:500 dilution; the p-AKT, HSP90, AKT, and Flag antibodies were prepared at a 1:1000 dilution. For cross-cell line western analysis of validation experiments, see **Supplementary** Fig. 1.

For western blot quantifications, ImageJ (version 2.9.0) was used to perform densitometry of PTEN expression relative to an HSP90 loading control across three replicates.^64^ The resulting values were then normalized to the average of the four controls included on each gel (C1, NT, U1, and D1). Two-sided pairwise-t-tests were then performed in R Studio (version 2022.12.0+353) using the R Stats Package (version 4.2.2) to assess the significance of the change in PTEN expression for each hit (H1-H13) relative the control group, which consisted of all four non-targeting (NT) or off-targeted controls (C1, U10, and D10).

### ABC Model Analyses

We implemented the ABC model (v0.2.2) as described.^16^ Inputs to the model included RCMC contact data and the following ENCODE data from the HCT116 cell line to determine candidate enhancer regions and quantify enhancer activity: DNAse-seq, H3K27Ac, and H3K4me1 (DNase-seq: ENCFF304MDQ, ENCFF969VWM; H3K27ac: ENCFF799ZUN, ENCFF977FPK; H3K4me1: ENCFF485QHQ, ENCFF119RZI). The promoter region was defined by the hg38 UCSC RefGene track (version 2017-03-08), centered on the start of the PTEN TSS. Within the model, we used the default Knight-Ruiz normalization and varied resolution (2KB, 5KB, 10KB, 15KB) when generating the normalized contact matrix and subsequent predictions.

### Statistics

Separately-processed biological samples were analyzed in biological triplicate at minimum for all experiments, and technical replicates within each biological replicate were performed as noted for individual experiments.

### Primers

All primers used in this study are listed in **Extended Data Table 4**.

### Antibodies

All antibodies used for staining or cell stimulation are listed in **Extended Data Table 5**.

### sgRNAs

All gRNAs used here are listed in **Supplementary Table 2**.

## Acknowledgements

This work was supported by Duke University School of Medicine start-up funds (K.C.W.), support from the Duke Cancer Institute (K.C.W.), Duke Office of Physician Scientist Development Strong Start support (C.E.E.), American Gastroenterological Association Research Scholar Award (C.E.E.), Massachusetts Institute of Technology Koch Institute support (Frontier Research Fund to A.S.H., Ludwig Center graduate fellowship to V.Y.G.), and NIH awards (R01CA207083 to K.C.W., K08CA263300 to C.E.E., F30CA247323 to C.G.C.S., K08CA259623 to S.E.J., DP2GM140938 and R33CA257878 to A.S.H and F32CA268527 to A.L.)

## Author Contributions

Conceptualization: CGCS, SEJ, CEE, KCW

Methodology: CGCS, HMH, VYG, ASH

Formal Analysis: CGCS, HMH, AL, VYG, CS, SCG, MJA

Investigation: CGCS, HMH, HK, KGB

Resources: VYG, CS, ASH, MJA

Writing – original draft: CGCS, CEE, KCW

Writing – review & editing: CGCS, HMH, KGB, CFB, SCG, ASH, SEJ, CEE, KCW

Visualization: CGCS, HMH, AL, VYG, CS

Funding acquisition: CEE, KCW

Supervision: CEE, KCW

Competing Interests:

## Additional Information

Supplementary Information is available for this paper in the “Extended Data” (Extended Data Figs. S1-S5, Extended Data Tables S1-S5) and Supplementary Files (Supplementary Fig. 1, Supplementary Table 1, and Supplementary Table 2)

## Correspondence and requests for materials

Correspondence and requests for materials should be addressed to C.E. Eyler and K.C. Wood.

## Data availability

Publicly available sequencing datasets utilized in the current work can be accessed using the accession numbers found in **Supplementary** Fig. 1). By the time of publication, all new sequencing data associated with our work will be made freely available via the Gene Expression Omnibus and/or Short Read Archive NCBI databases.

