## Extended Data for "Integrative *PTEN* Enhancer Discovery Reveals a New Model of Enhancer Organization"

**The PDF file includes:**

Extended Data Figs. 1 to 5
Extended Data Table 1 to 5

**Other Supplementary Materials for this manuscript include the following:**

Supplementary Fig. 1
Supplementary Table 1 and 2

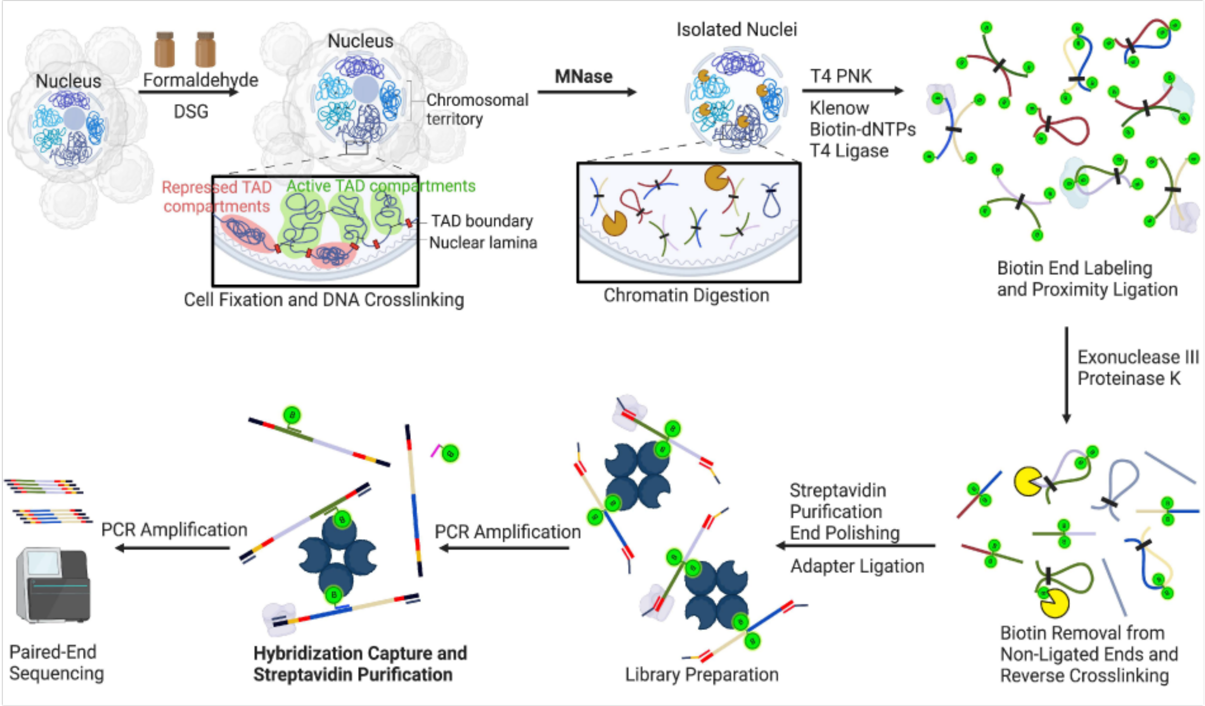


**Extended Data Fig. 1. RCMC achieves nucleosome resolution by combining micrococcal nuclease fragmentation, locus-specific oligo capture, and efficient deep sequencing.** Schematic overview of the RCMC protocol. HCT116 cells were crosslinked with DSG and formaldehyde. The chromatin was digested to mononucleosomes utilizing a micrococcal nuclease (MNase) digestion, followed by end-repair with biotin labeling, and proximity ligation. Samples were then reverse-crosslinked, size-selected for dinucleosome fragments, and purified for ligated fragments via streptavidin pulldown. The resultant DNA was processed into Micro-C sequencing libraries via adapter ligation and PCR. We designed a custom 120-mer oligo capture pool tiling a 4.16Mb window surrounding the *PTEN* locus; the capture pool was designed at 2x coverage of the genomic window with filtering for highly repetitive regions, ultimately yielding a pool of 60,570 oligos that covered 90.16% of the region. The capture pool was synthesized and purchased as a Custom Target Enrichment Panel from Twist Bioscience. Oligo-mediated locus-specific hybridization capture and RCMC sequencing library preparation were then performed using our custom oligo capture pool following the Twist Fast Hybridization Target Enrichment Protocol. The library was sequenced on the NovaSeq 6000 platform (Illumina).

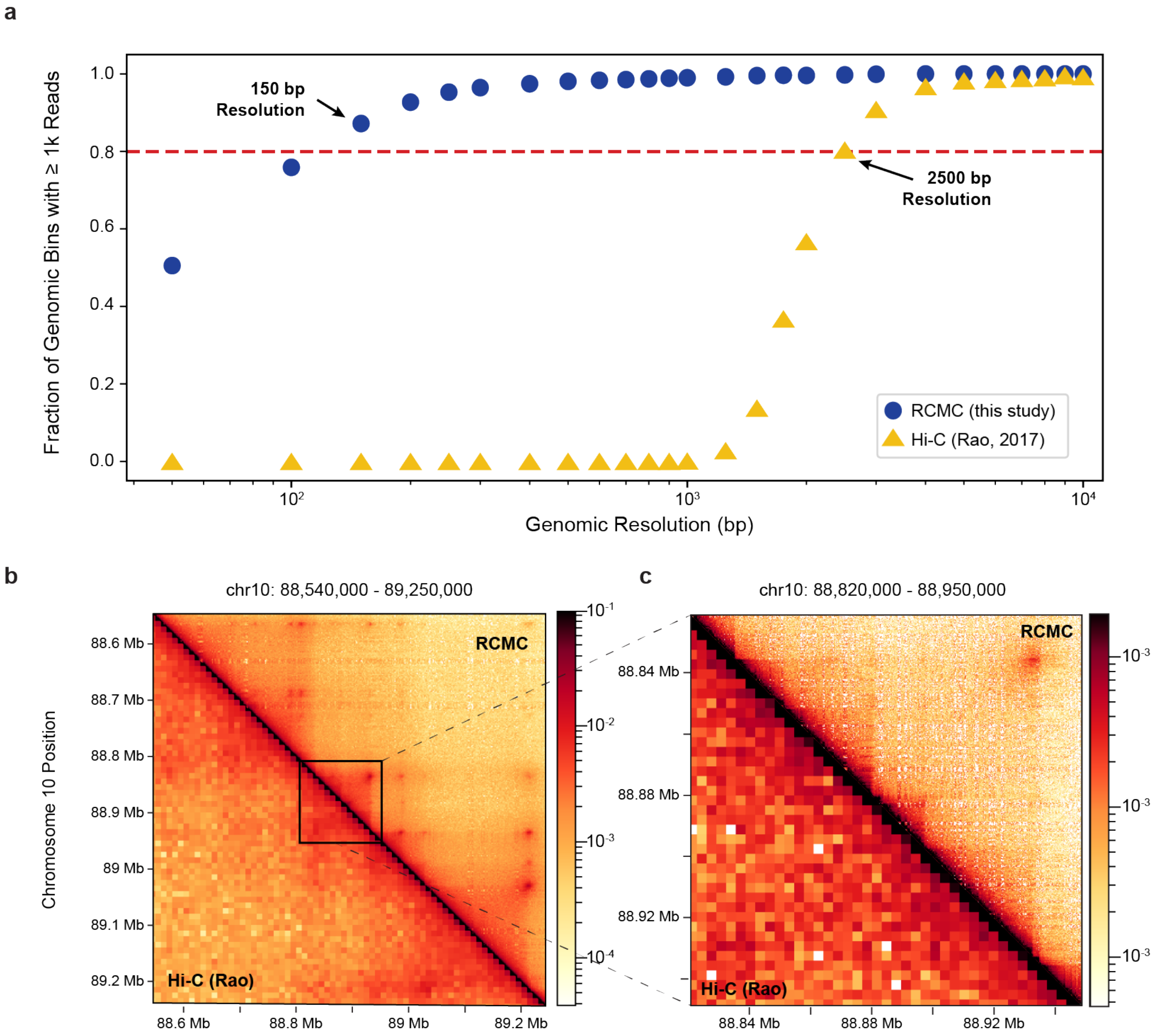


**Extended Data Fig. 2. RCMC yields a dramatically enhanced nucleosome-resolution contact map for a large genomic window enabling detection of novel structure. (a)** Calculation of contact map resolution for RCMC versus Hi-C (Rao et al, 2017). Plot depicts the percentage of bins with greater than 1000 reads as a function of bin size. Resolution is determined by the bin size for which greater than 80% of bins contain at least 1000 reads. Blue circles=RCMC (this study), yellow triangles=Hi-C, red dashed Line=80% of bins containing at least 1000 reads. Contact map comparison of RCMC against the previously deepest available HCT116 Hi-C dataset scaled to **(b)** a 740 kb window (visually optimized with RCMC bin size=2 kb, Hi-C bin size=10 kb), and zoomed-in to **(c)** a 130 kb window (visually optimized with RCMC bin size=250bp and Hi-C bin size=3200bp, both approaching calculated maximum resolutions).

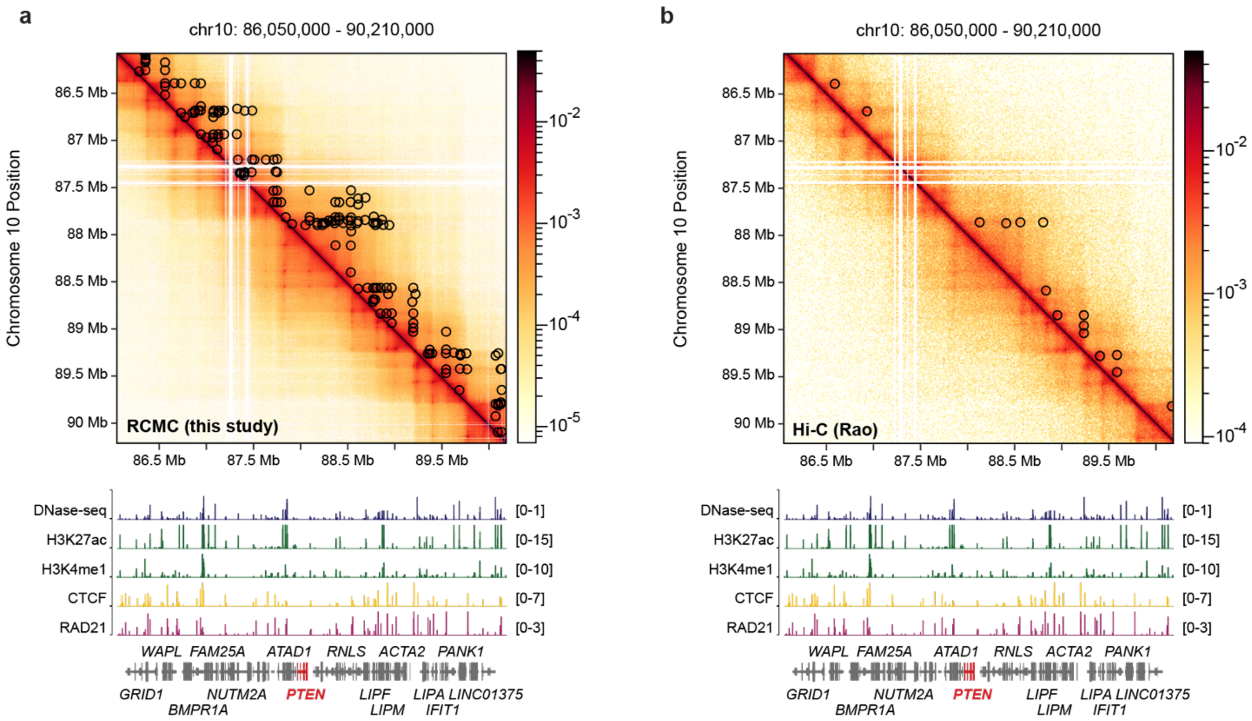


**Extended Data Fig. 3. Enhanced resolution enabled by RCMC yields improved loop calling at the *PTEN* locus.** Contact map showing loop calls from RCMC study across the *PTEN* capture window of 4.16Mb for **(a)**. our RCMC study, versus **(b)** previously deepest available HCT116 Hi-C dataset (Rao et al, 2017). For panels **(a)** – **(b)**: visualization of each contact map was optimized to the technique and viewing window; here, RCMC=2kb bins and Hi-C=10kb bins. Loop interactions called by Mustache algorithm are shown as black circles on each upper reflection. Gene annotations are shown, *PTEN* is in red. DNase-seq=dark blue, H3K27ac=green, H3K4me1=green, CTCF=yellow and RAD21=pink.

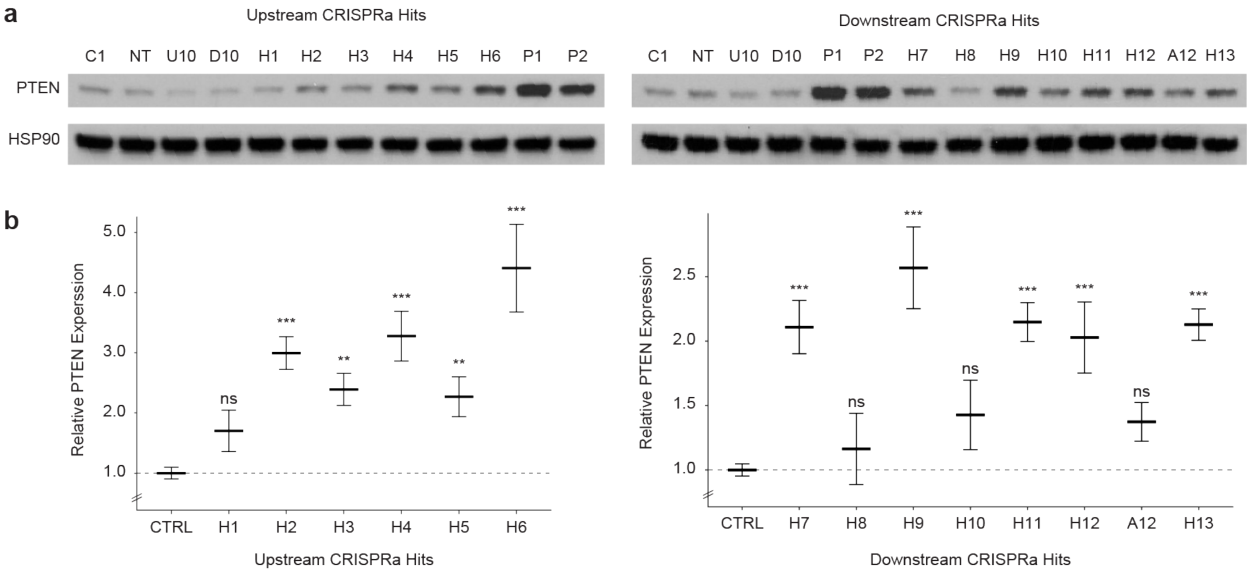


**Extended Data Fig. 4. Protein-level validation of tiled CRISPRa screen-nominated *PTEN* enhancers (a) Single sgRNA validation of all CRISPRa-nominated hit regions relative to control via western blot in HCT116.** NT=non-targeting control, Chr1=negative control targeting Chr1, U10=negative control targeting upstream region on Chr10, D10= negative control targeting downstream region on Chr10, P1=promoter positive control 1, P2=promoter positive control 2, A12=RCMC anchor nearest H12. (b) Densitometry of PTEN expression relative to HSP90 loading control for single sgRNA validations of all CRISPRa-nominated hit regions, n=3, mean and standard error depicted in plot. CTRL=compilation of negative controls (C1, NT, U10, and D10). Pairwise-t-tests were performed to determine statistical significance relative to CTRL; ns=not significant, *=p<0.05, **=p<0.01, ***p<0.001.

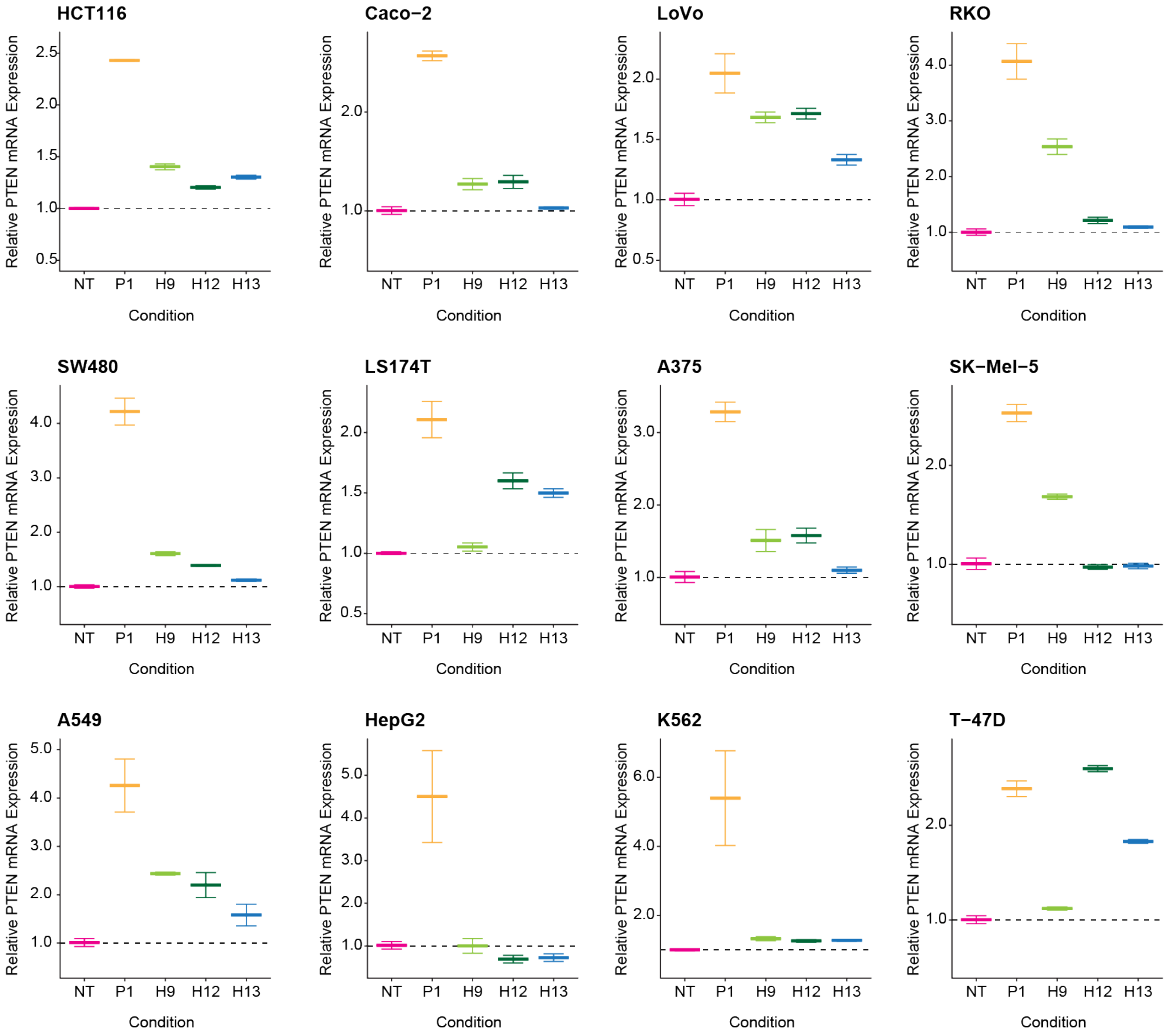


**Extended Data Fig. 5. mRNA-level validation of tiled CRISPRa screen-nominated *PTEN* enhancers across cell lines.** Single sgRNA validation of select CRISPRa-nominated hit regions ("H9", "H12", and "H13") relative to controls (NT=nontargeting control, P1=promoter positive control) via qRT-PCR across cell line panel. Mean and standard error depicted here for n=3 biological replicates assayed in quadruplicate technical replicates for PTEN and internal normalization control.

#### Extended Data Table 1.
#### Significantly scoring regions in functional CRISPRa PTEN FACS screen.

| Hit ID | Chr10 Start | Chr10 End | Hit Midpoint on Chr10 | Genomic Relationship to PTEN | Composite DHS Region | Distance from PTEN TSS | Previously Described as PTEN Enhancer | Hit Region Max Deconvolved Score (CRISPR-SURF) | Hit Region Min Adjusted p-value (CRISPR-SURF) | Functionally Validated by Western Blot |
|---|---|---|---|---|---|---|---|---|---|---|
| H1 | 87817762 | 87818342 | 87818052 | Upstream | 282 | -45573 | No | 0.017298 | 0.0001 | No |
| H2 | 87842382 | 87842502 | 87842442 | Upstream | 285 | -21183 | No | 0.051231 | 0.0001 | Yes |
| H3 | 87843102 | 87843382 | 87843242 | Upstream | 286 | -20383 | No | 0.042648 | 0.0001 | Yes |
| H4 | 87846062 | 87846302 | 87846182 | Upstream | 289 | -17443 | No | 0.073168 | 0.0001 | Yes |
| H5 | 87850802 | 87851082 | 87850942 | Upstream | 291 | -12683 | No | 0.030685 | 0.0001 | Yes |
| H6 | 87860162 | 87860482 | 87860322 | Upstream | 292 | -3303 | No | 0.08401 | 0.0001 | Yes |
| P1 | 87861542 | 87863082 | 87862312 | Promoter | 293 | -1313 | Yes (Kwon et al 2022) | 0.083875 | 0.0001 | Yes |
| P2 | 87863342 | 87863722 | 87863532 | Promoter | 293 | -93 | No | 0.140747 | 0.0001 | Yes |
| P3 | 87863742 | 87864462 | 87864102 | Promoter | 293/294 | 477 | No | 0.073783 | 0.0001 | Did not test (redundant promoter) |
| P4 | 87865782 | 87866082 | 87865932 | Promoter | 294 | 2307 | No | 0.037527 | 0.0001 | Did not test (redundant promoter) |
| H7 | 88028602 | 88028942 | 88028772 | Downstream | 311 | 165147 | No | 0.03233 | 0.0001 | Yes |
| H8 | 88108362 | 88108602 | 88108482 | Downstream | 322 | 244857 | No | 0.025706 | 0.0001 | No |
| H9 | 88110342 | 88110702 | 88110522 | Downstream | 323 | 246897 | No | 0.0859 | 0.0001 | Yes |
| H10 | 88114522 | 88114642 | 88114582 | Downstream | 324 | 250957 | No | 0.026447 | 0.0001 | No |
| H11 | 88254182 | 88254382 | 88254282 | Downstream | 345 | 390657 | No | 0.016212 | 0.01 | Yes |
| H12 | 88385442 | 88385782 | 88385612 | Downstream | 362 | 521987 | Yes (Tottone et al 2020) | 0.055664 | 0.0001 | Yes |
| H13 | 88428082 | 88429182 | 88428632 | Downstream | 370 | 565007 | No | 0.051523 | 0.0001 | Yes |

#### Extended Data Table 2.
#### Cell lines and culture conditions.

| Cell Line | Cell Culture Medium |
|---|---|
| A375 | RPMI 1640, 10% FBS |
| A549 | F-12K, 10% FBS |
| Caco-2 | MEM, pyruvate, NEAA, 20% FBS |
| HCT116 | McCoy’s 5A Medium, 10% FBS |
| HEK293T | DMEM (High Glucose), NEAA, Sodium Pyruvate, GlutaMAX, 10% FBS |
| HepG2 | McCoy’s 5A Medium, 10% FBS |
| K562 | IMDM, 10% FBS |
| LoVo | F-12K, 10% FBS |
| LS174T | MEM, pyruvate, NEAA, 10% FBS |
| RKO | MEM, pyruvate, NEAA, 10% FBS |
| SK-Mel-5 | MEM, pyruvate, NEAA, 10% FBS |
| SW480 | RPMI 1640, 10% FBS |
| T-47D | RPMI 1640, pyruvate, HEPES, glucose, 10% FBS , 0.2 IU/mL insulin (35 µL of 4 mg/mL in 20 mL) |

**Extended Data Table 3.**
**Region Capture Micro-C (RCMC) versus Hi-C mapping comparison.**

| | Whole dataset | | | | | | |
|---|---|---|---|---|---|---|---|
| | Total sequencing reads | Mapped sequencing reads | | Mapped sequencing reads (unique) | | Unique contacts, >= 1 kb | |
| | | Read count | % total | Read count | % mapped | Read count | % uniquely mapped |
| **Region Capture Micro-C (RCMC)** | 791,968,270 | 437,772,998 | 55.3% | 208,163,049 | 47.6% | 158,487,430 | 76.1% |
| **Hi-C (Rao et al., 2017)** | 3,475,834,764 | 1,435,303,924 | 41.3% | 1,310,830,680 | 91.3% | 940,803,149 | 71.8% |

| | ***PTEN* locus** (*hg38* chr10:86,050,000-90,210,000; locus size = 4.16 Mb) | | | | | |
|---|---|---|---|---|---|---|
| | Mapped sequencing reads | | Mapped sequencing reads (unique) | | Unique contacts, >= 1 kb | |
| | Read count | % all mapped | Read count | % mapped | Read count | % uniquely mapped |
| **Region Capture Micro-C (RCMC)** | 295,150,512 | 67.4% | 88,071,236 | 29.8% | 73,515,231 | 83.5% |
| **Hi-C (Rao et al., 2017)** | 3,298,767 | 0.2% | 3,009,199 | 91.2% | 2,468,921 | 82.0% |

### Extended Data Table 4.
### PCR primers and TaqMan assays.

| **CRISPRa Screen Custom PCR Primers** | |
| --- | --- |
| Fwd 5' | AATGATACGGCGACCACCGAGATCTACACAATTTCTTGGGTAGTTTGCAGTT |
| Rev 5' | CAAGCAGAAGACGGCATACGAGAT-(6 bp index sequence)-GACTCGGTGCCACTTTTTCAA |

| **CRISPRa Screen Sequencing Primers** | |
| --- | --- |
| Custom Read 5' | GATTTCTTGGCTTTATATATCTTGTGGAAAGGACGAAACACCG |
| Custom Index 5' | GCTAGTCCGTTATCAACTTGAAAAAGTGGCACCGAGTC |

| **Puro Amplification Primers** | |
| --- | --- |
| PCR PuroR Primers FWD | TGGAGGAGAATCCCGGCCCTGGTACCATGGTGTCTAGAGAGTACAAGCCCACGGTG |
| PCR PuroR Primers rev | TTGTAAGTCATTGGTCTTAAAGGTACCTGAGGTGTGACTGGAAAACCCACCTCCTC |

| Gene Expression Assay | Company (catalogue number) | Probe ID |
| --- | --- | --- |
| PTEN | Thermofisher (4351370) | Taqman Probe PTEN: Hs02621230_s1 FAM-MGB |
| TBP | Thermofisher (4331182) | Taqman Probe TBP: Hs00427620_m1 VIC-MGB |

**Extended Data Table 5.**
**Antibodies (western and flow activated cell sorting).**

| Western Blots | Antibody Name | Company |
| --- | --- | --- |
| PTEN | Purified Mouse Anti-PTEN | BD Biosciences |
| phospho-AKT | Phospho-Akt (Ser473) | Cell Signaling Technology |
| HSP90 | HSP90 (C45G5) Rabbit mAb | Cell Signaling Technology |
| AKT | Akt (pan) (C67E7) Rabbit mAb | Cell Signaling Technology |
| Flag | Monoclonal ANTI-FLAG® M2 antibody | Sigma |

| PTEN Flow Screen | Antibody Name | Company |
| --- | --- | --- |
| PTEN | Alexa Fluor® 647 Mouse anti-PTEN | BD Biosciences |
