## Supplementary Fig 1 for "Integrative *PTEN* Enhancer Discovery Reveals a New Model of Enhancer Organization"

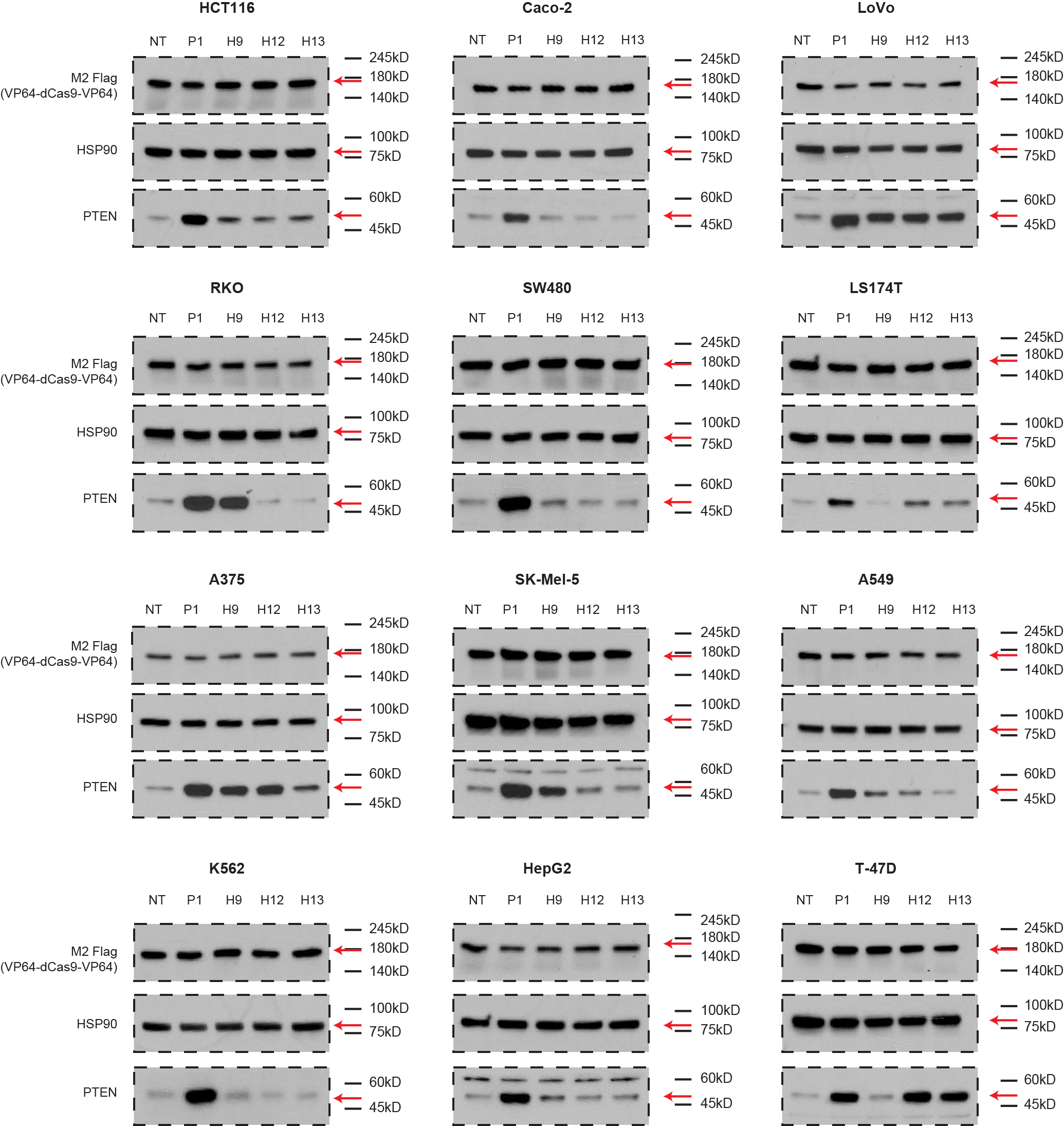

Supplementary Fig. 1. Validation of tiled CRISPRa screen-nominated *PTEN* enhancers across 12 cell lines. Single sgRNA validation of select CRISPRa-nominated hit regions (“H9”, “H12”, and “H13”) relative to controls (NT=nontargeting control, P1=promoter positive control) via western blot across cell line panel.
